# Individual Prediction of Psychotherapy Outcome in Posttraumatic Stress Disorder using Neuroimaging Data

**DOI:** 10.1101/647925

**Authors:** Paul Zhutovsky, Rajat M. Thomas, Miranda Olff, Sanne J.H. van Rooij, Mitzy Kennis, Guido A. van Wingen, Elbert Geuze

## Abstract

**Objective:** Trauma-focused psychotherapy is the first-line treatment for posttraumatic stress disorder (PTSD) but 30-50% of patients do not benefit sufficiently. We investigated whether structural and resting-state functional magnetic resonance imaging (MRI/rs-fMRI) data could distinguish between treatment responders and non-responders on the group and individual level.

**Methods:** Forty-four male veterans with PTSD underwent baseline scanning followed by trauma-focused psychotherapy. Voxel-wise gray matter volumes were extracted from the structural MRI data and resting-state networks (RSNs) were calculated from rs-fMRI data using independent component analysis. Data were used to detect differences between responders and non-responders on the group level using permutation testing, and the single-subject level using Gaussian process classification with cross-validation.

**Results:** A RSN centered on the bilateral superior frontal gyrus differed between responders and non-responder groups (*P*_FWE_ < 0.05) while a RSN centered on the pre-supplementary motor area distinguished between responders and non-responders on an individual-level with 81.4% accuracy (*P* < 0.001, 84.8% sensitivity, 78% specificity and AUC of 0.93). No significant single-subject classification or group differences were observed for gray matter volume.

**Conclusions:** This proof-of-concept study demonstrates the feasibility of using rs-fMRI to develop neuroimaging biomarkers for treatment response, which could enable personalized treatment of patients with PTSD.

## Introduction

Posttraumatic stress disorder (PTSD) is a psychiatric disorder which can develop after experiencing a traumatic event. It is characterized by states of re-experiencing of the traumatic event, avoidance of trauma-reminders, emotional numbing, and hyperarousal (1). PTSD lifetime prevalence rates in the general population are estimated to be below 10% (varying between 1.3% to 8.8% depending on the country) (2) but can vary heavily in veterans (between 1.4% to 31%) (3, 4). Treatment of PTSD typically involves trauma-focused psychotherapy with or without the administration of medication such as selective serotonin reuptake inhibitors (SSRIs). Trauma-focused therapies such as trauma-focused cognitive behavior therapy (TF-CBT) or eye movement desensitization and reprocessing (EMDR) have been suggested as first-line treatments for treating PTSD (5). However, 30-50% of patients do not benefit sufficiently (6). To improve treatment response rates it is important to better understand differences between responders and non-responders, and identify reliable predictors for treatment outcome.

PTSD is characterized as a brain disorder showing differences in activity and connectivity of large-scale brain networks (7). The connectivity of these networks can be recorded using neuroimaging techniques such as resting-state functional magnetic resonance imaging (rs-fMRI). Therefore, it is important to investigate if those alterations in rs-fMRI connectivity could be used to predict treatment-outcome and reveal biomarkers to increase the treatment-response rate. Indeed, pre-treatment group differences in fMRI activity and connectivity were observed between responders and non-responders in PTSD in several studies (8-11). However, these group-level univariate analyses focus on average differences between responders and non-responders. This does not allow inference at the individual patient level, which can be achieved using multivariate supervised machine learning analyses (12). Most importantly, performance is evaluated on new data to estimate the generalizability of the trained models, and thereby enabling the prediction of treatment outcome for new patients. Machine learning analyses have been performed in the context of PTSD using different modalities of MRI data to distinguish between patients and controls (13, 14). However, only two studies to date have used machine learning analyses to predict future outcome at an individual level. One study aimed to predict clinical status two years after treatment with 12 weeks of paroxetine in a sample of 20 civilian PTSD patients (15). This study used pre- and post-treatment rs-fMRI derived measures, amplitude of low frequency fluctuations and whole-brain degree centrality maps, and the results showed that pre-but not post-treatment measures were able to predict remission status after two years with an accuracy of 72.5%. But as all but one patient had been in remission shortly after treatment, these results reflect relapse rather than treatment outcome. In addition, one recent study used a combination of resting-state connectivity within the ventral attention network and delayed recall performance in a verbal memory task to predict the response to prolonged exposure therapy in ∼19 civilian patients with PTSD (16). Although the proportion of treatment non-responders was low, the classifier still managed to distinguish the groups with ≥80% sensitivity and specificity.

To determine whether neuroimaging data could also predict treatment outcome in a larger sample of combat veterans with PTSD, we analyzed pre-treatment structural MRI and rs-fMRI data of 44 patients who received treatment-as-usual. This consisted of trauma-focused psychotherapy such as TF-CBT and EMDR, and clinical outcome was determined 6-8 month following the baseline fMRI scan. We previously reported pre-treatment group differences in structural (17), white-matter (18) and task-based (f)MRI (8, 19) between responders and non-responders, as well as rs-fMRI differences between patients and controls (20) and post-treatment rs-fMRI differences between responders and non-responders (21). For the present study, we derived maps of regional gray matter volume using voxel-based morphometry (VBM). In addition, we extracted functional connectivity (FC) within resting-state networks (RSNs) using independent component analysis (ICA). To ensure independence between RSN identification and estimation of RSN expression for each individual patient, the ICA was performed on rs-fMRI data of sex and age matched combat controls (n=28). Subsequently, we performed univariate inference on the group level as well as multivariate prediction on the single-subject level using Gaussian process classification (GPC) with 10×10 cross-validation.

## Methods

### Participants

In total 57 veterans with PTSD and 29 combat controls (CC) were included in the study. Patients were recruited from one of four outpatient clinics of the Military Mental Healthcare Organization in Utrecht, The Netherlands. PTSD diagnosis was established by a licensed psychologist or psychiatrist. The Clinician Administered PTSD scale (CAPS) (22) for DSM-IV (1) was administered by trained research staff to quantify the total symptom severity and had to be ≥45. Combat controls had to have no current psychiatric disorders and a total CAPS score <15. Further inclusion criteria for all subjects were deployment to a war zone and 18-60 years of age. Comorbid disorders were examined using the structured clinical interview for DSM-IV (SCID-I) (23). Subjects with a history of neurological disorders, current substance dependence and contraindications for MRI scanning were excluded. From the initial 57 PTSD patients, seven were lost to follow-up, three were excluded based on excessive motion during scanning (see Supplemental Methods section), one due to an artifact in the MRI scan, and one due to refusal of scanning. One additional participant was excluded as she was the only female in the sample. This leads to the final sample of 44 PTSD patients. From the CCs only one subject had to be excluded based on excessive motion (n = 28).

After a period of six to eight months in which patients underwent treatment-as-usual consisting of trauma-focused therapy (e.g. TF-CBT, EMDR) a second CAPS assessment was performed. Treatment response was defined as a ≥30% decrease of total CAPS score at follow-up with respect to the baseline assessment (24, 25). According to this criterion 24 PTSD veterans were defined as responder and 20 as non-responder. All participants gave written informed consent. The study was approved by the University Medical Center Utrecht ethics committee, in accordance with the declaration of Helsinki (26).

### Clinical Data Analysis

To estimate whether the CCs, responders and non-responders differed across any demographic or clinical variables at baseline or follow-up ANOVA, Kruskal-Wallis, χ^2^, or t-tests were applied as appropriate. All tests were performed using the R software (version 3.5.1).

### Data Acquisition

All scans were obtained on a 3T MRI scanner (Philips Medical System, Best, the Netherlands). The T1-weighted high resolution MRI scan was acquired before the rs-fMRI scan with the following parameters: repetition time (TR) = 10ms, echo time (TE) = 4.6ms, flip angle = 8°, 200 sagittal slices, field of view (FOV) = 240 × 240 × 160, matrix size = 304 × 299 and voxel size = 0.8 × 0.8 × 0.8mm. The rs-fMRI scan consisted of 320 T2*-weighted echo planar interleaved slices with TR = 1600ms, TE = 23ms, flip angle = 72.5°, FOV = 256 × 208 × 120, 30 transverse slices, matrix size = 64 × 51, total scan time 8 min and 44.8 s, 0.4 mm gap, acquired voxel size = 4 × 4 × 3.60mm). Participants were asked to focus on a fixation cross, while letting their mind wander and relax.

### MRI data preprocessing

To estimate whether structural images carry information to distinguish between responders or non-responders a VBM analysis was performed. Gray matter (GM) voxel-wise volume maps were computed using the SPM12 toolbox (v7219, https://www.fil.ion.ucl.ac.uk/spm/software/spm12/). Resting-state fMRI images were preprocessed using the advanced normalization tools (ANTs, 2.1.0, http://stnava.github.io/ANTs/) and FMRIB Software Library (FSL, 5.0.10, https://fsl.fmrib.ox.ac.uk/fsl/fslwiki/. To control for the influence of motion on the rs-fMRI data ICA-AROMA was applied (27). Details on the preprocessing pipelines can be found in the Supplemental Methods section.

### Resting State Network Identification

Preprocessed rs-fMRI data were analyzed to determine group-level resting-state networks (RSNs). Group components with a fixed number of 70 components were estimated using a meta-ICA approach utilizing FSL’s MELODIC software (28) applied to the rs-fMRI data of the CCs. We chose to only use the CCs in this step to ensure that the definition of the RSNs and the machine learning analysis were independent from each other. The meta-ICA approach allows for the identification of reproducible and reliable group components (29). After meta-ICA, 48 RSN’s were identified using a semi-automatic approach. Thereafter, FSL’s dual regression approach was used to estimate single-subject spatial representations of the corresponding group networks for all patients. Details on the implementation and rationale of the procedure can be found in the Supplemental Methods section, and signal and noise components are illustrated in Supplemental Figure 1 and Supplemental Figure 2.

### Univariate Analysis

The preprocessed GM volume maps from the VBM analysis and the identified RSNs were used to investigate group differences between responders and non-responders. Age and total intracranial volume were entered as covariates for the VBM data, while only age was used as covariate for the RSN data. The significance level was set to *P* < 0.05 family-wise error (FWE) corrected and estimated using the threshold-free-cluster-enhancement statistic (TFCE) (30) with permutation testing (10000 permutations) using the TFCE toolbox (r167, http://dbm.neuro.uni-jena.de/tfce/) for the VBM data. For the resting-state data, the PALM toolbox (a112, https://fsl.fmrib.ox.ac.uk/fsl/fslwiki/PALM) was used, since it allowed for permutation-based FWE correction across the whole-brain and all 48 RSNs at the same time. Both analyses accounted for two-tailed tests.

### Multivariate Analysis

For the multivariate single-subject classification of responders and non-responders, we used the GM volume-maps from the VBM analysis and each RSN separately. Classification was performed using a Gaussian process classifier (GPC) (31). Briefly, GPCs are multivariate Bayesian classifiers which allow to obtain valid probabilistic predictions by estimating the posterior distribution, given a pre-defined prior distribution. Univariate feature selection was performed on the training set to reduce the initial data dimension using nested 5-fold cross-validation (see Supplemental Methods). Performance was estimated by calculating sensitivity, specificity, balanced accuracy, area under the receiver-operator curve (AUC) and positive/negative predictive value (PPV/NPV) using ten times repeated 10-fold cross-validation to avoid overfitting bias. To estimate whether our classifier performed better than chance, label permutation tests with 1000 iterations were performed. The final *P*-values were Bonferroni corrected for 49 tests.

We also investigated the performance of the GPC when an uncertainty option was allowed: utilizing the probabilistic output of the classifier, we established regions of uncertainty for which the classifier would not make a prediction. For example, with an uncertainty region of 10% any probabilistic GPC output for a new patient which lies between 45-55%, would not be assigned a classification label (because the classification into responders and non-responders would be uncertain). Only patients with a higher (or lower) probability would be assigned to a class and considered for calculation of balanced accuracy. This allowed us to investigate how well our GPC would perform if classification has only to be made if a specific level of certainty is reached and how many patients would need to be excluded to reach that level.

## Results

### Clinical data

Demographic information, clinical variables and outcomes of statistical tests can be found in Table 1. There was no difference in demographics between the CCs, responders or non-responders, nor any clinical difference between responders and non-responders at baseline. At follow-up non-responders showed a higher total CAPS score (t(42) = 7.89, *P* < 0.001) and higher use of serotonin reuptake inhibitors (χ^2^(1) = 5.77, *P* = 0.02).

**Table 1:**
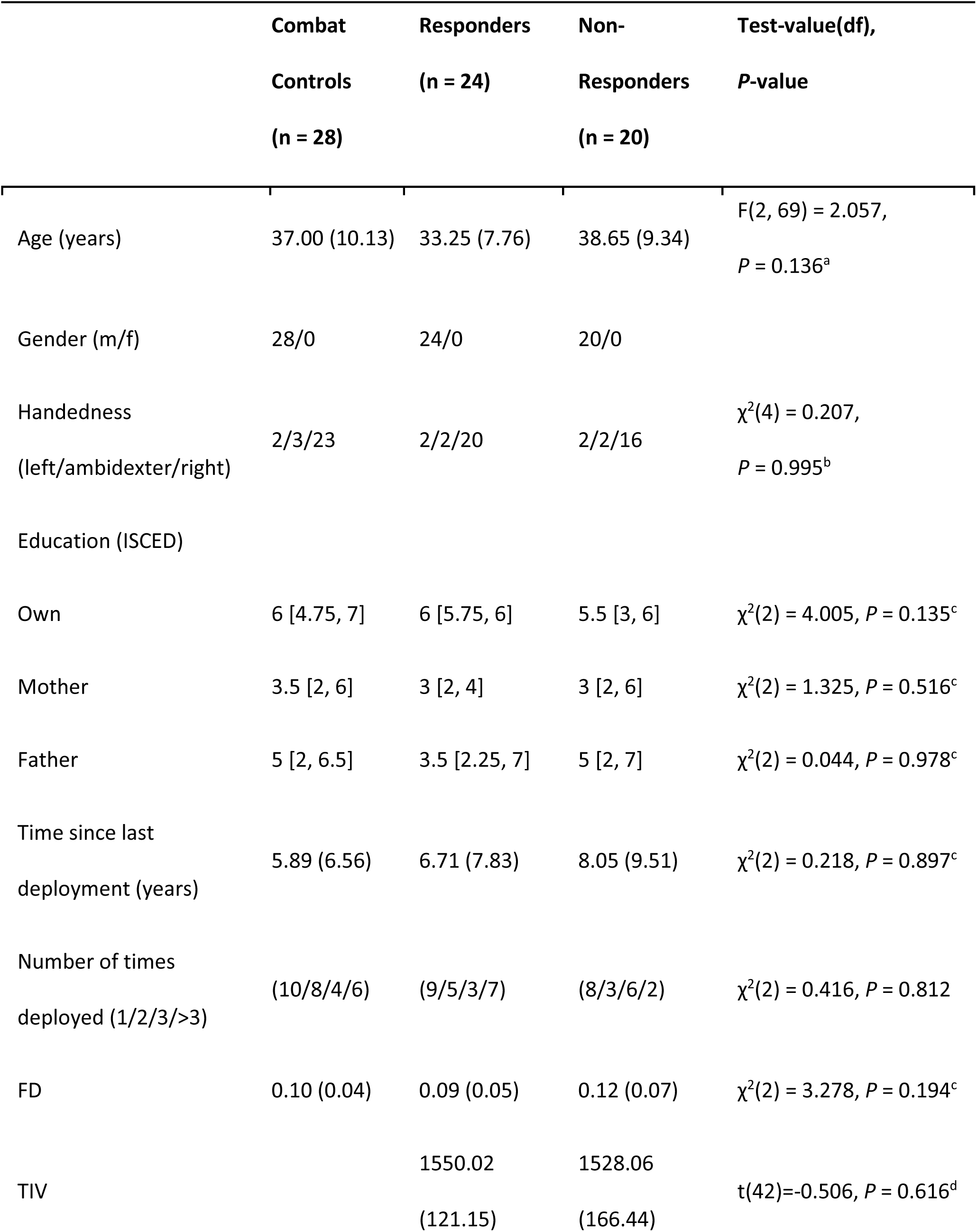
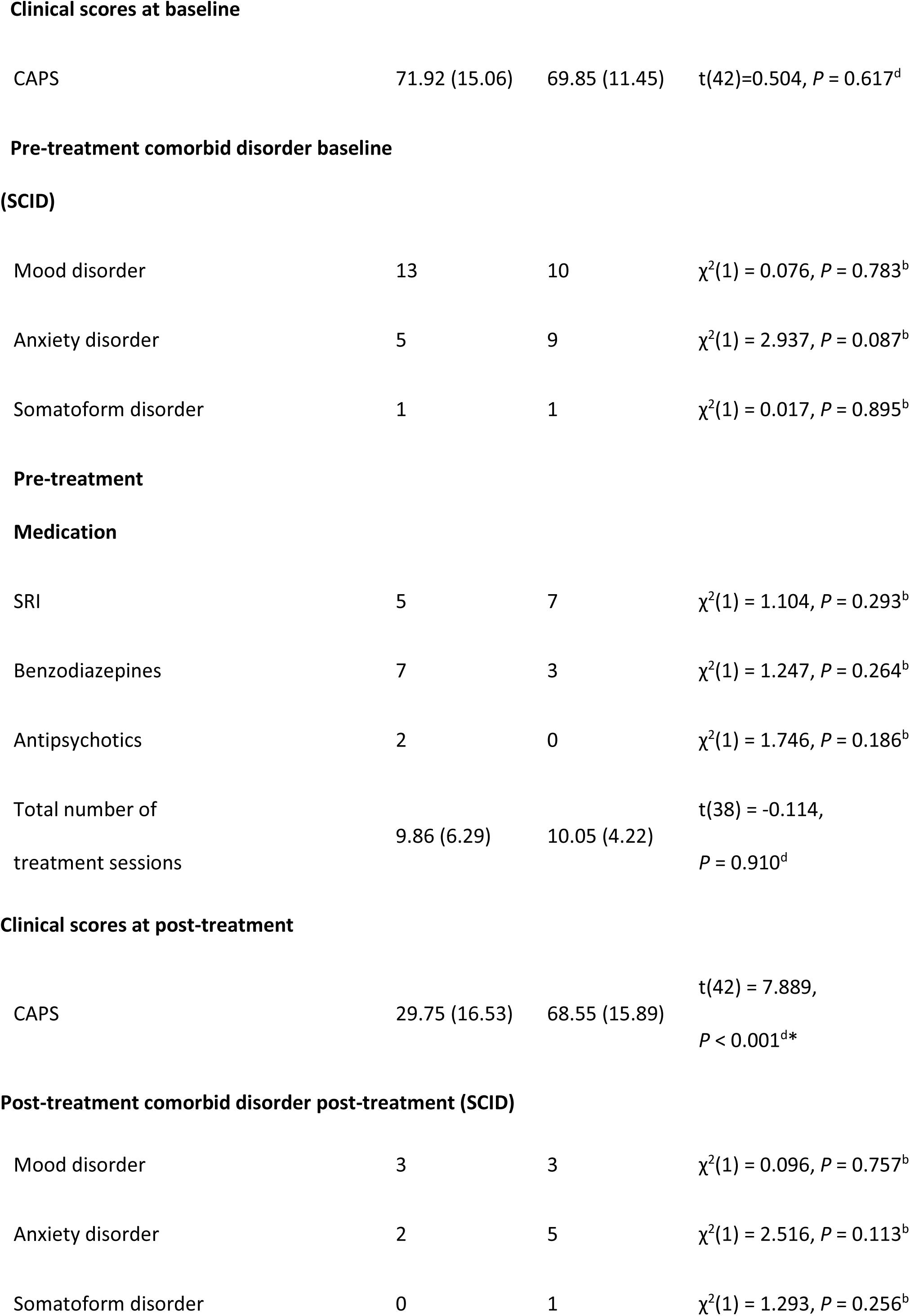
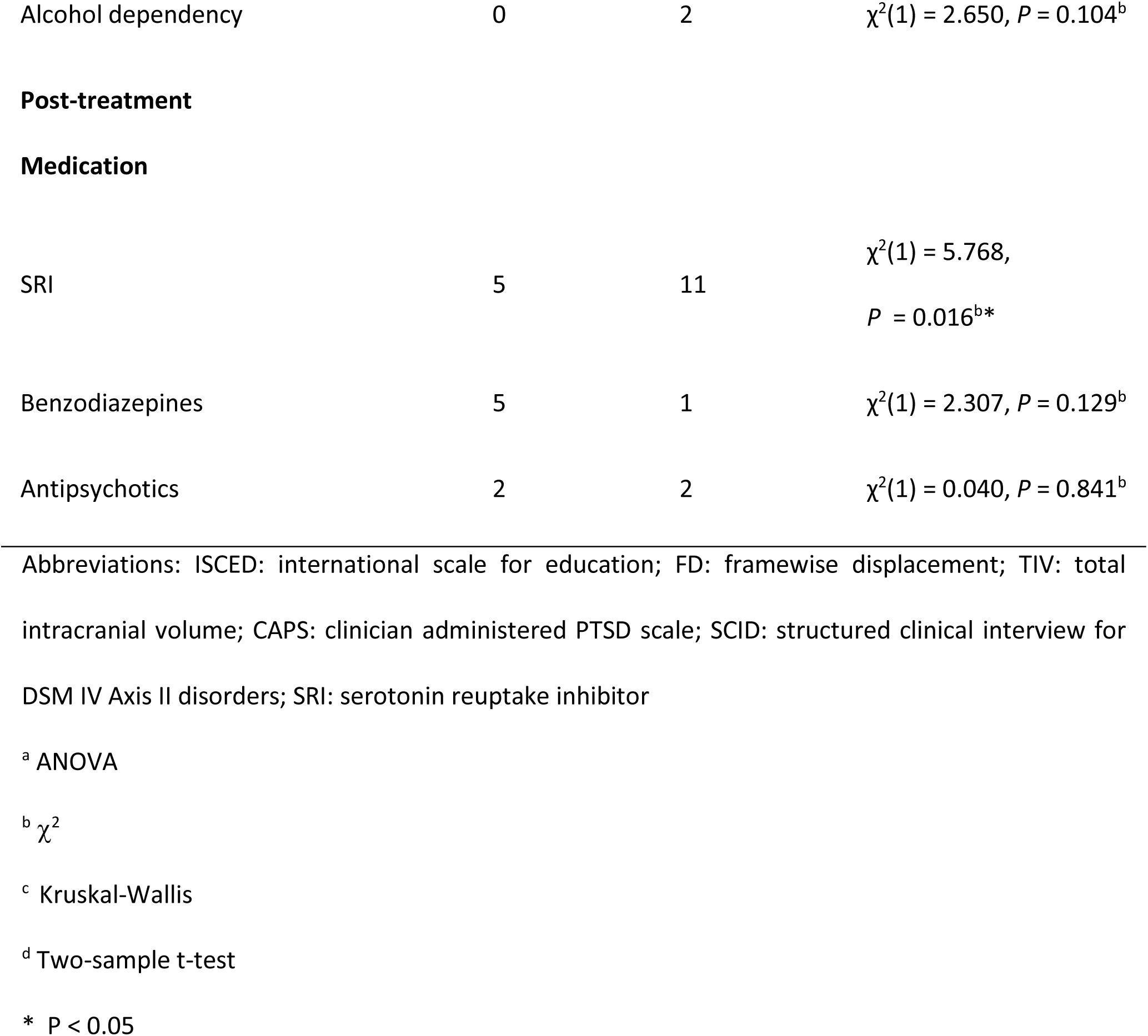
Demographics and clinical data

### Univariate analysis

After correction for multiple comparisons across all RSNs, the rs-fMRI analysis showed one network with significantly increased connectivity in non-responders as compared to responders (Figure 1). The network was centered on the bilateral lateral frontal polar area and the difference was observed in the right superior frontal gyrus (*P*_FWE_ = 0.04). No significant group differences in GM were observed.

**Figure 1.**
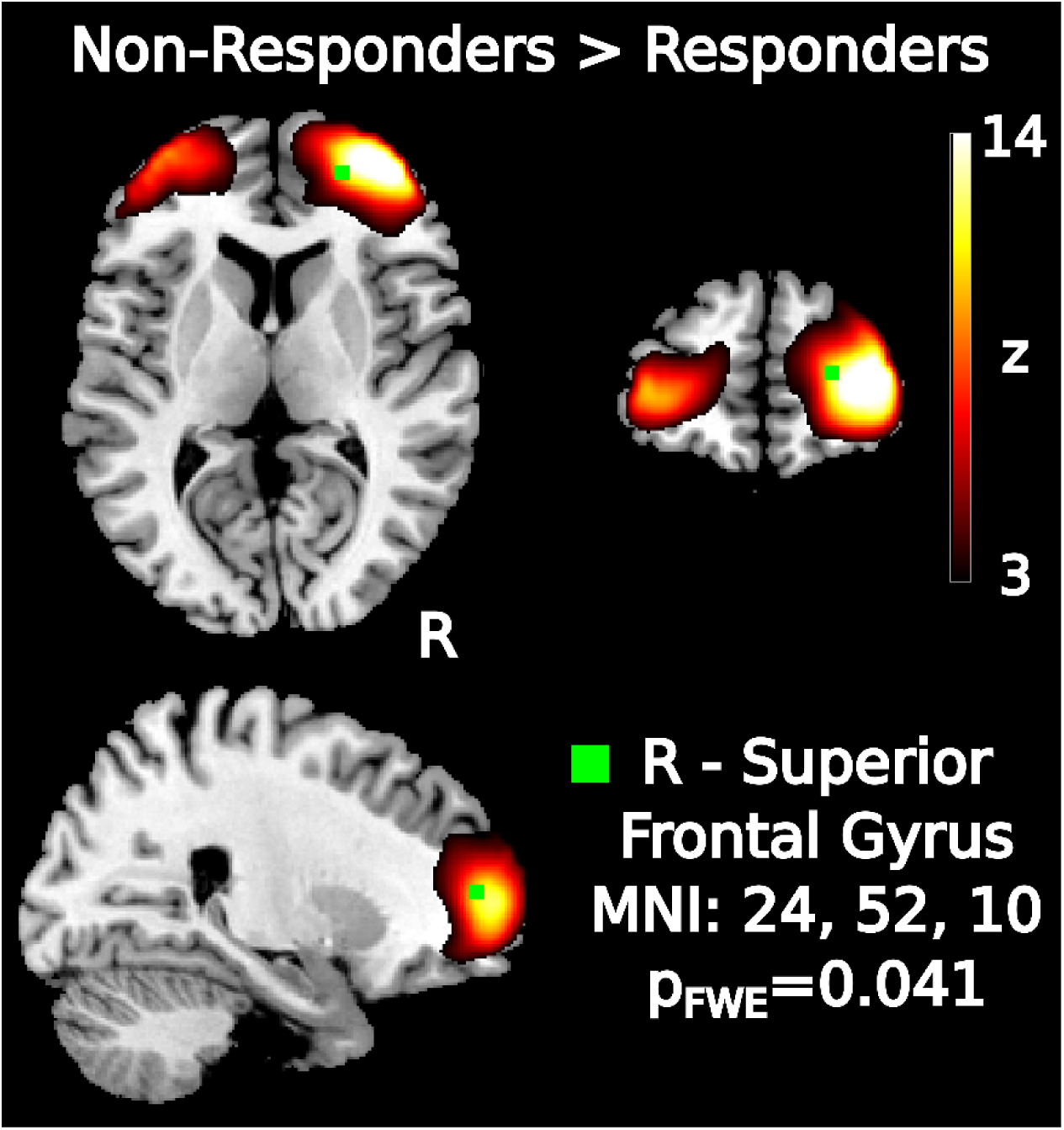
Results of the group-level univariate RSN analysis. Higher resting-state connectivity in non-responders than responders in the frontopolar network. Two-tailed *P*-value was corrected for whole-brain comparisons and 48 networks.

### Multivariate analysis

GPC’s trained on a network centered around the pre-supplementary motor area (pre-SMA) could classify non-responders and responders with an average cross-validated balanced accuracy of 81.4% (SD: 17.2, *P*_Bonferroni_ < 0.05) (Figure 2A). The network showed excellent AUC (0.929, SD: 0.149) with high sensitivity (84.8%, SD: 25.1), moderately high specificity (78% SD: 28.6), and high PPV/NPV (0.840/0.835, SD: 0.214/0.262). No other network showed significant classification performance after Bonferroni correction was applied, including the network that showed a significant difference on the group-level in the univariate analysis.

**Figure 2.**
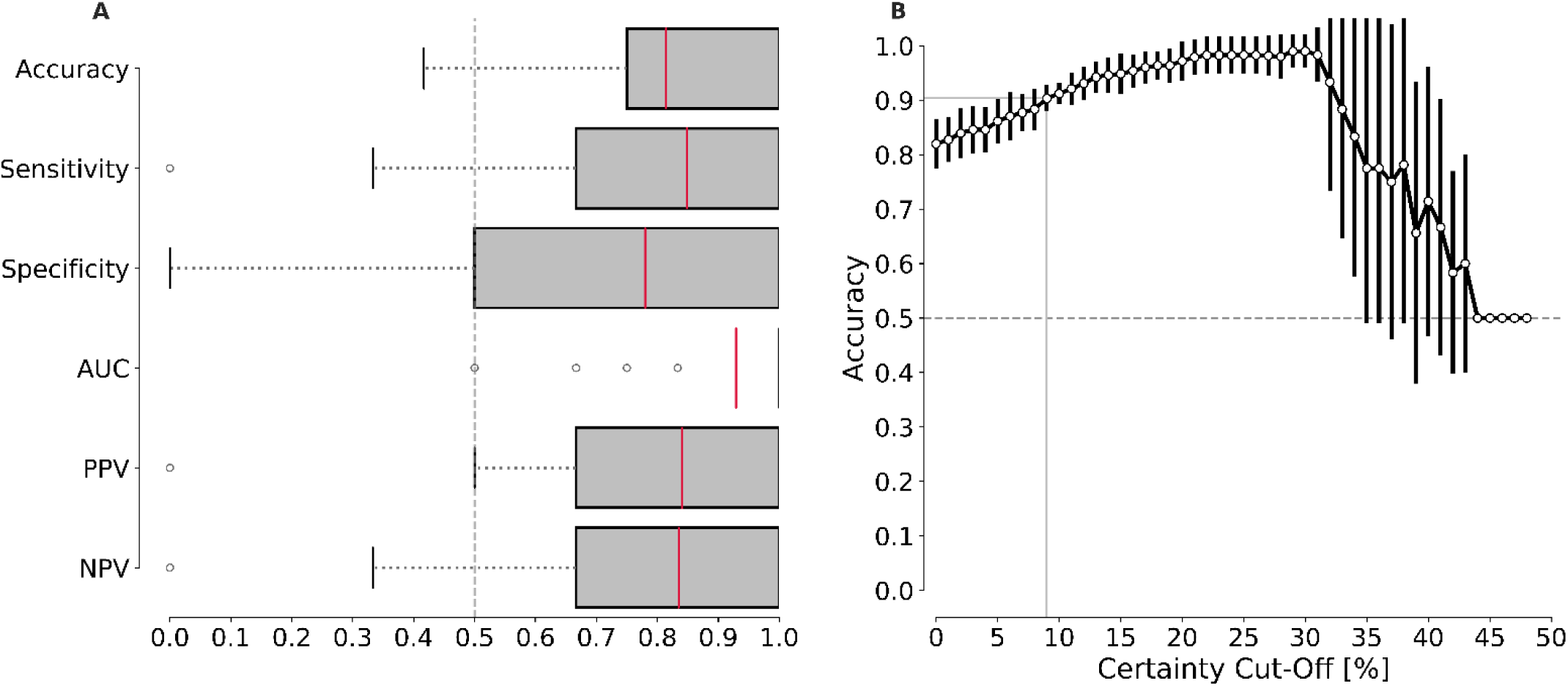
Results of the single-subject multivariate prediction analysis of treatment outcome. A) Classification metrics of the pre-SMA network shown as box-and-whisker plots. Outliers plotted as circles were determined as values which lay outside 1.5 times the interquartile range. Please note that the box for the AUC metric collapsed because the first quartile and the median were the same value. B) Post-hoc evaluation of accuracy of GPC classifier when a certain level of certainty has to be ensured averaged across the ten repetitions of the 10-fold cross-validation with SD plotted as error bars. For example, once 12 patients (27%) with low prediction certainty of 0.41-0.59 – where 0.5 is equal probability of prediction – would be excluded, accuracy would increase to over 90%.

To investigate which regions of the pre-SMA network were most important for the classification process we examined consistently selected voxels during the feature selection process. We tracked the selection frequency of voxels across cross-validation runs, looking at voxels which were selected in >50% of the runs (Table 2 and Figure 3). Regions in both hemispheres located outside the group-network were contributing to the classification performance. The largest clusters were located in the left inferior temporal gyrus (n_voxel_ = 14), left superior frontal gyrus (n_voxel_ = 10), and right precentral gyrus (n_voxel_ = 9).

**Table 2:**
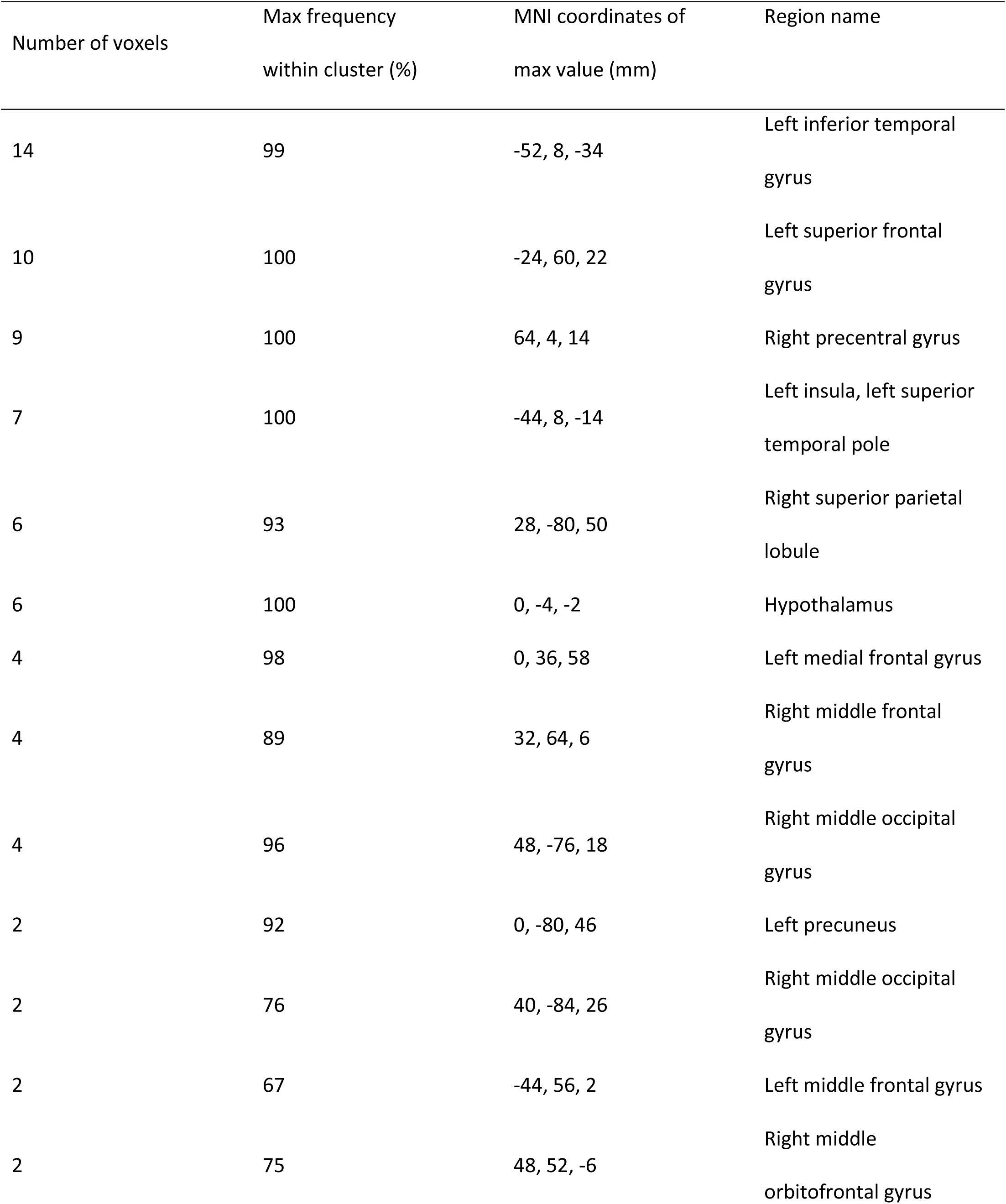
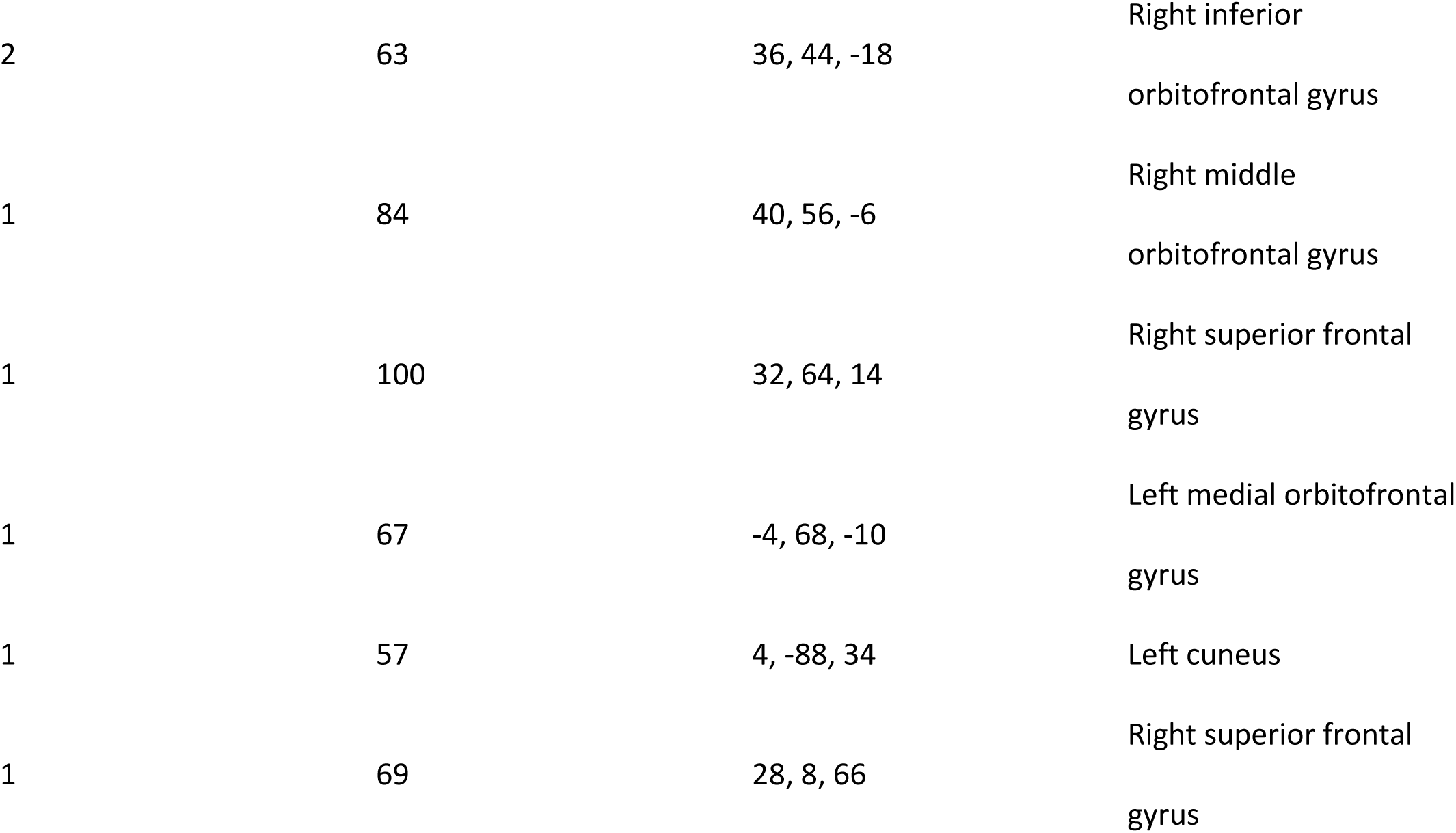
Most frequently selected features during the nested-cross-validation procedure of the pre-SMA network

**Figure 3.**
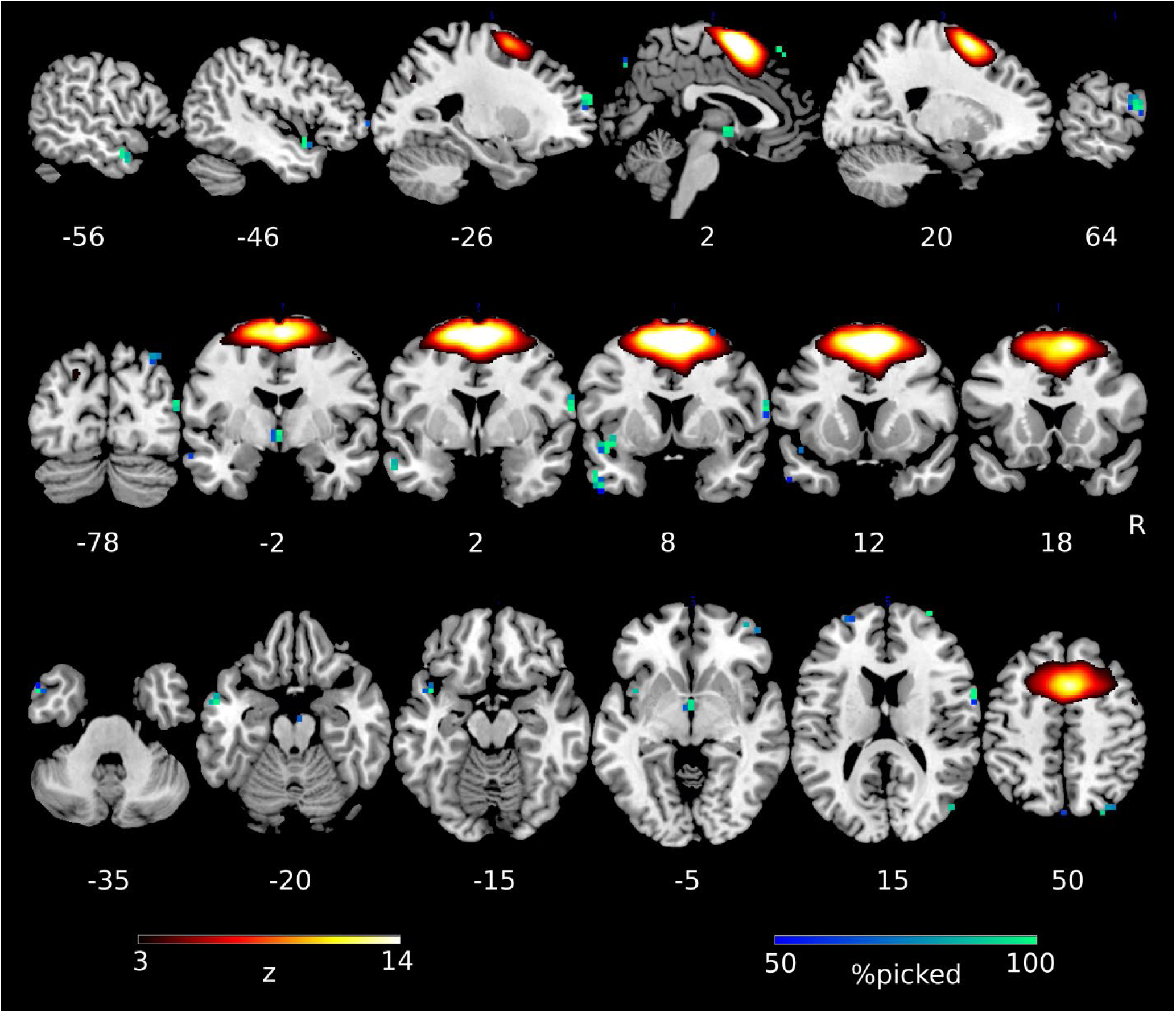
Best performing network in the multivariate classification (pre-SMA) in hot colors and the most often selected voxels during the classification in cold colors.

Additionally, we provided a post-hoc evaluation of what would happen if prediction would only be made for patients for which a high degree of certainty of the classifier is established. As illustrated in Figure 2B, this ability to ‘reject’ patients from the classification with increasing classification certainty leads to increasing accuracy while at the same time reducing the number of patients for which the GPC can make a classification. For example, once 12 patients (27%) with low prediction certainty of 0.41-0.59 – where 0.5 is equal probability of prediction – would be excluded, accuracy would increase to over 90%.

## Discussion

The present study investigated the possibility of using pre-treatment structural MRI and rs-fMRI data to predict the response to trauma-focused psychotherapy in male combat veterans with PTSD. The results showed that rs-fMRI data successfully distinguished between responders and non-responders univariate and multivariate analyses. The univariate analysis detected group differences in a network centered on the frontal pole, and the multivariate analysis predicted treatment response on an individual level using pre-SMA connectivity with an accuracy of 81.4%. Whereas previous studies have focused on MRI-based treatment outcome predictors at the group level, our results suggest that single-subject prediction is also feasible. This result provides a proof-of-concept for the feasibility of developing predictive biomarkers, which could enable personalized treatment for patients with PTSD.

Our multivariate analysis revealed the predictive importance of the pre-SMA. This brain area is closely linked to the SMA and both areas are involved in motor execution and imagination. However, the pre-SMA can also be distinguished from the SMA, and is more involved in higher cognitive processes such as working memory, language and task switching (32, 33). Furthermore, the pre-SMA has been implicated to be involved in the process of response inhibition which has been previously related to PTSD development and treatment-response in PTSD (see (34) for a review). Therefore, the discovered network might relate to the level of cognitive control in these patients, however, future studies are needed to elucidate why individual differences within this network are important for PTSD trauma-focused therapy response. Intriguingly, resting-state connectivity within this network is also predictive for the response to electroconvulsive therapy in depression (35). The main difference in results is that the network in the current study is more confined to the pre-SMA due to the use of ICA with 70 components instead of 32 components, which was associated with a larger network that consisted of a large part of the dorsomedial prefrontal cortex. Together, this suggests that pre-SMA connectivity may determine responsiveness to treatment, regardless of intervention and disorder.

The discovered network is different from the ventral attention network (VAN, consisting of the insula, dorsal anterior cingulate, anterior middle frontal gyrus, and supramarginal gyrus) that was recently reported. The VAN in combination with delayed recall performance in a verbal memory task could predict prolonged exposure therapy outcome in a sample of ∼19 civilians with PTSD with sensitivity and specificity ≥80% (16). But even though both studies used rs-fMRI, the underlying biomarkers cannot readily be compared. First, the variables tested in (16) were discovered by performing comparisons between healthy controls and PTSD patients, whereas we discovered the pre-SMA network from comparisons between responders and non-responders directly. Second, the authors did not investigate any other networks beyond the VAN for treatment outcome prediction. And third, the brain regions that are part of the VAN were actually part of distinct RSNs in our ICA analysis, whereas the VAN was considered one network in the other study. Therefore, it remains to be tested whether VAN or pre-SMA connectivity is also predictive in other samples. Regardless, both studies demonstrate that rs-fMRI contains information that is informative for predicting psychotherapy outcome on an individual level.

The univariate group analysis showed increased connectivity in non-responders in the frontal pole. The frontal pole region (BA 10) has been implicated in a multitude of cognitive tasks, including attention, perception, language, and memory tasks (36, 37). Specifically, the lateral parts of the frontal pole are more associated with working memory and episodic memory retrieval while medial parts of the frontal pole were mostly involved in mentalizing, which is the reflection of your own emotions and mental states (36, 37). This division of the frontal pole was recently confirmed by a cytoarchitectonic parcellation indicating two distinct areas: a more lateral frontal pole area 1 (FP 1) and a more medial frontal pole area 2 (FP 2). Our frontal polar network was mostly located in FP 1 and may therefore be associated with memory related processes. Interestingly, the frontal pole is particularly known for its role in metacognitive processes such as prospective memory, which refers to the ability to remember to perform an intended action in the future (38). Prospective memory allows maintaining and retrieving future goals and plans, which is expected to be relevant for the success of psychotherapy. Additionally, a proposed underlying mechanism of psychotherapy action is memory reconsolidation, which refers to the process of modifying maladapted memories (39). We therefore speculate that the observed difference in the FP 1 region might reflect one or both of these mechanisms.

The difference between the identified networks in the univariate and multivariate analyses might seem counterintuitive at first but can be explained by the differences in objective and methodology of both analyses. This discrepancy is in line with the observation that significant group-level differences do not necessarily translate to high classification accuracies because of strongly overlapping distributions and different goals of the analysis (12). A significant *P*-value in a group-level analysis does not have to correspond to the ability of distinguishing between individual patients because the statistically significant difference in average values might show low effect sizes. In these case classification performance will be low. In addition, the goal of statistical inference is the identification of localized differences between groups while the goal of classification is to find the best multivariate combination of data which would allow to generalize the effect to new subjects. These are two inherently different goals which therefore can lead to different outcomes.

In contrast to our results, previous studies that have used univariate analysis of structural MRI and task-based fMRI data have primarily pointed to pre-treatment differences in the anterior cingulate cortex, amygdala, hippocampus and insula (8-11). However, direct comparison with our study is difficult since these differences might be due to the use of task-based fMRI, usage of a predefined region of interest approach, different types of psychotherapy, different PTSD populations and different criteria for treatment response (40). This can be exemplified with the absence of results for the structural MRI analysis which is in contrast to our previous finding of differences in hippocampal volume between patients with remitted vs. persistent PTSD (17). This difference could be due to the calculation of the volumes: in the present study, a VBM analysis was employed to provide a highly multivariate data set which could be optimally used during the classification procedure, whereas we previously estimated hippocampal volume using segmentation in Freesurfer. In addition, in this study we chose to focus on treatment response while previously we investigated the more stringent criterion of treatment remission to focus on PTSD persistence.

The current study has several limitations. Although the study is among the largest for treatment outcome prediction using neuroimaging in psychiatry, the sample size is small for machine learning analyses with many more features than subjects. This could result in high variance of the estimated accuracy and the results therefore require further validation in independent samples. Another limitation of this study is the use of an all-male veteran sample. This limits the generalization of the results to other patients with PTSD. Therefore, a replication of the proposed approach in a more diverse sample would be desirable. Finally, the treatments received by the patients represent a heterogeneous mix of different trauma-focused psychotherapies. While they are considered as first-line treatments and the fact that in realistic settings multiple treatments might be employed by psychiatrists, the results are not specific to one particular treatment. This can also be considered a strength, as the predictive power generalizes to different types of psychotherapy.

In conclusion, the current study shows that treatment response to trauma-focused psychotherapy can be predicted for individual patients with PTSD using machine learning analysis of rs-fMRI data. This proof-of-concept study demonstrates the feasibility to develop neuroimaging biomarkers for treatment response, which will enhance personalized treatment of patients with PTSD.

## Supplementary Material

### Supplemental Methods

#### MRI processing

The voxel-based morphometry (VBM) analysis was performed using the SPM12 toolbox. Briefly, we obtained gray matter (GM) segmentations applying the unified segmentation approach (41). The GM maps were then normalized to MNI space (1.5mm3) based on DARTEL registration (42) using a template derived from 555 healthy controls of the IXI-database (http://www.brain-development.org) in MNI space provided by the CAT12 toolbox (http://www.neuro.uni-jena.de/cat/). The normalized GM images were modulated by the Jacobian determinant to preserve local tissue volume and spatially smoothed with a kernel of 8mm at FWHM.

#### fMRI processing

Preprocessing of fMRI images was performed using the advanced normalization tools (ANTs, 2.1.0, http://stnava.github.io/ANTs/) (43) and FMRIB Software Library (FSL, 5.0.10) (44). For the purpose of registration to MNI space and extraction of white matter (WM) and cerebral spinal fluid (CSF) signal from fMRI scans, T1 images were bias-field corrected using the N4 algorithm (45) and brain-extracted using scripts from the ANTs toolbox and the Oasis template (https://www.oasis-brains.org/). Images were then segmented into GM, WM and CSF partial volume estimates using FSL’s FAST (46). The skull-stripped images were normalized to MNI space using the ANTs symmetric normalization procedure (43). One PTSD patient was excluded based of an artifact in his MRI scan.

fMRI image preprocessing consisted of realignment, co-registration to the T1 image using boundary-based registration (47) and spatial smoothing with a 8mm FWHM kernel. Motion has a strong effect on resting-state fMRI measures (48) and therefore has to be addressed further during the preprocessing of rs-fMRI data. Therefore, we calculated framewise displacement (FD) (49) of the raw data and excluded subjects based on the following criteria: (1) any rotation/translation parameter >4mm/°,(2) average FD > 0.45,(3) m ore than 150 volum es with an individual FD of 0.25, leading to less than 4min of motion free rs-fMRI data (50).

Applying these criteria led to the exclusion of three PTSD patients and one combat control. The remaining patients did not differ in their motion levels (see Table 1). Furthermore, motion was additionally addressed by applying ICA-AROMA (27) to automatically identify single-subject ICA components associated with motion. These components were then regressed out from the data. Further structured noise was removed by performing nuisance regression with average WM and CSF signals. For that the calculated WM/CSF segmentations of the T1 image were transformed to EPI space and thresholded conservatively at 0.95. The denoised fMRI images were transformed to MNI space at 4mm and high-pass filtered at 0.01Hz.

#### Meta-ICA

For the meta-ICA we repeatedly (25 times) extracted 20 participants out of the 28 combat controls at random and performed a temporally-concatenated group-ICA with the number of components fixed to 70. The obtained spatial maps (25 * 70 = 1750) were merged and entered into an additional (meta-)ICA with 70 components. The number of components was determined because it was shown to provide good insight into clinical differences of patient groups (51). Following the meta-ICA, group spatial components were investigated visually and using an automatic approach verifying their reproducibility across individual ICA runs and the proportion of the components located in the gray matter (52). 48 components were identified as carrying non-noise related resting-state activity (Supplementary Figure 1 and Supplementary Figure 2). Following the identification of the components dual regression was performed to identify subject specific spatial maps corresponding to the group components (53). Dual regression was applied using the group maps computed through meta-ICA on the combat controls and the rs-fMRI data of the PTSD patients.

#### Multivariate Analysis

Classification was performed using a Gaussian process classifier (GPC). We chose a zero mean and a normalized linear kernel function for the prior distribution following recommendations in the field (54). To infer the parameters of the posterior distribution we used a Probit likelihood function with the expectation maximization algorithm for inference (55). To reduce the initial dimensionality of the classification problem univariate feature selection was performed. For that we computed the average univariate difference between connectivity values in every voxel using only participants of the training set. The difference was then z-scaled and thresholded. To determine the optimal threshold we investigated z-values from 2.5 until 4.0 in steps of 0.1 using nested 5-fold cross-validation on the training set. The optimal value was chosen as the one which generated the highest average balanced accuracy (average between sensitivity and specificity) across the five folds. The GPC was implemented using the Python (version 2.7.15) interface of the Shogun machine learning toolbox (version 6.1.3, http://shogun-toolbox.org/).

**Supplementary Figure 1.**
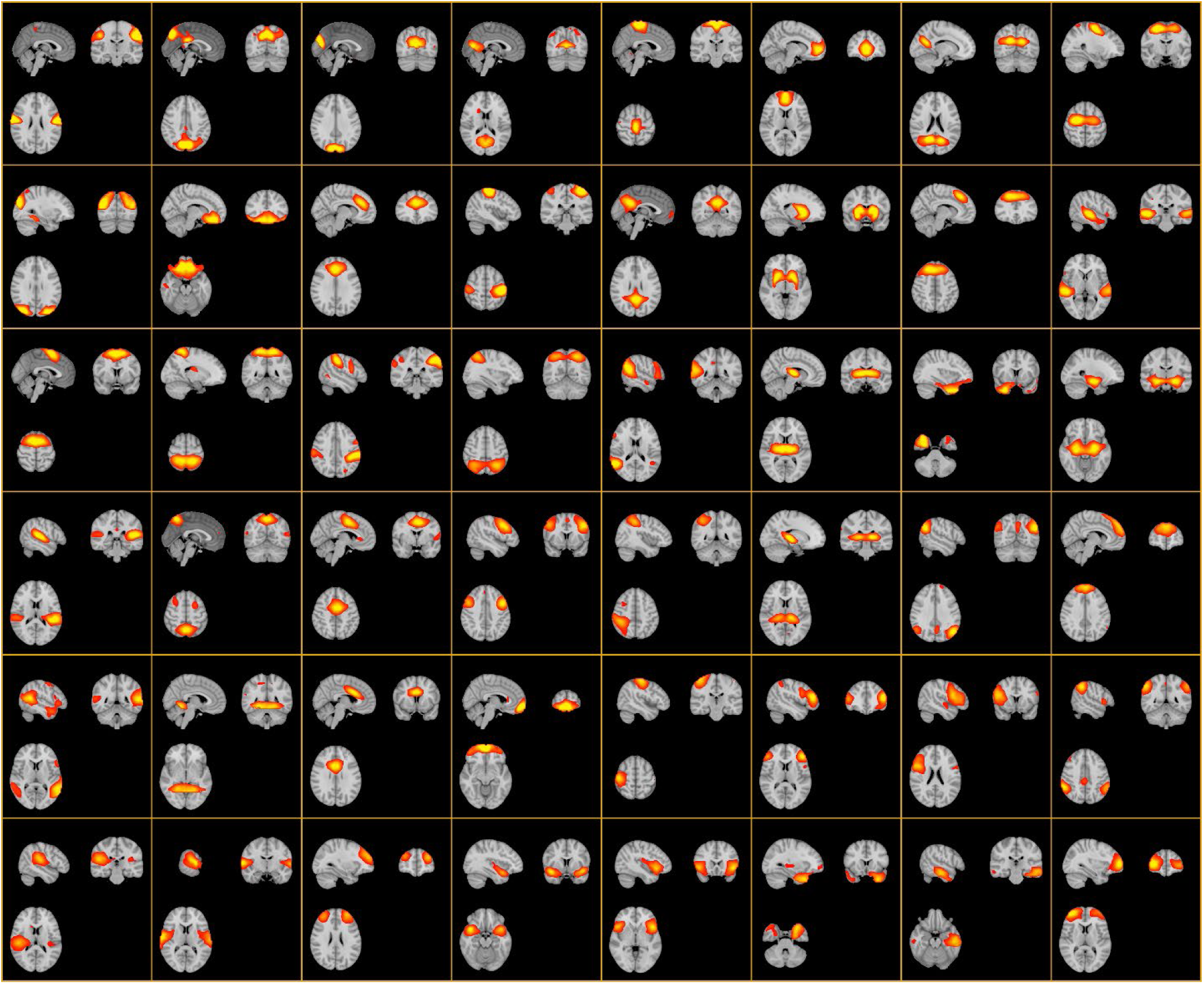
48 out of 70 networks calculated via a meta-ICA approach which were identified as carrying signal-related information

**Supplementary Figure 2.**
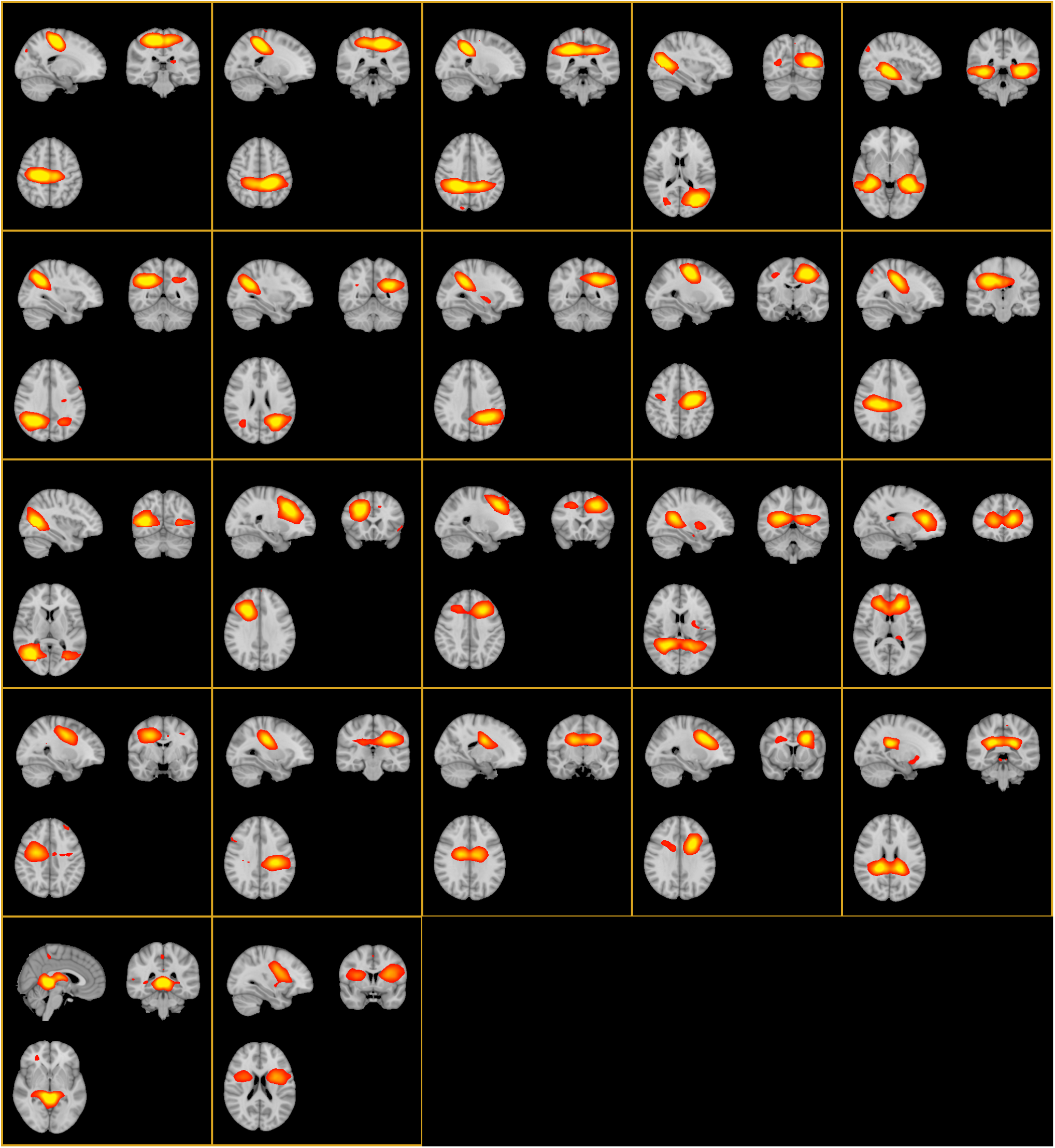
22 out of 70 meta-ICA networks which were related to noise sources.

## Acknowledgments

This study was supported by the Dutch Ministry of Defense, the Netherlands Organization for Scientific Research (NWO/ZonMW Vidi 016.156.318) and the AMC Research Council (150622).

## References

1. American Psychiatric A: Diagnostic and statistical manual of mental disorders: DSM-IV. Washington, DC, American Psychiatric Association; 1994.

2. Atwoli L, Stein DJ, Koenen KC, McLaughlin KA. Epidemiology of posttraumatic stress disorder: prevalence, correlates and consequences. Curr Opin Psychiatry. 2015;28:307–311.

3. Richardson LK, Frueh BC, Acierno R. Prevalence estimates of combat-related post-traumatic stress disorder: critical review. Aust N Z J Psychiatry. 2010;44:4–19.

4. Sundin J, Fear NT, Iversen A, Rona RJ, Wessely S. PTSD after deployment to Iraq: conflicting rates, conflicting claims. Psychol Med. 2010;40:367–382.

5. Cusack K, Jonas DE, Forneris CA, Wines C, Sonis J, Middleton JC, Feltner C, Brownley KA, Olmsted KR, Greenblatt A, Weil A, Gaynes BN. Psychological treatments for adults with posttraumatic stress disorder: A systematic review and meta-analysis. Clin Psychol Rev. 2016;43:128–141.

6. Bradley R, Greene J, Russ E, Dutra L, Westen D. A multidimensional meta-analysis of psychotherapy for PTSD. Am J Psychiatry. 2005;162:214–227.

7. Fenster RJ, Lebois LAM, Ressler KJ, Suh J. Brain circuit dysfunction in post-traumatic stress disorder: from mouse to man. Nat Rev Neurosci. 2018;19:535–551.

8. van Rooij SJ, Geuze E, Kennis M, Rademaker AR, Vink M. Neural correlates of inhibition and contextual cue processing related to treatment response in PTSD. Neuropsychopharmacology. 2015;40:667–675.

9. Aupperle RL, Allard CB, Simmons AN, Flagan T, Thorp SR, Norman SB, Paulus MP, Stein MB. Neural responses during emotional processing before and after cognitive trauma therapy for battered women. Psychiatry Res. 2013;214:48–55.

10. Falconer E, Allen A, Felmingham KL, Williams LM, Bryant RA. Inhibitory neural activity predicts response to cognitive-behavioral therapy for posttraumatic stress disorder. J Clin Psychiatry. 2013;74:895–901.

11. Fonzo GA, Goodkind MS, Oathes DJ, Zaiko YV, Harvey M, Peng KK, Weiss ME, Thompson AL, Zack SE, Lindley SE, Arnow BA, Jo B, Gross JJ, Rothbaum BO, Etkin A. PTSD Psychotherapy Outcome Predicted by Brain Activation During Emotional Reactivity and Regulation. Am J Psychiatry. 2017;174:1163–1174.

12. Bzdok D, Ioannidis JPA. Exploration, Inference, and Prediction in Neuroscience and Biomedicine. Trends Neurosci. 2019.

13. Gong Q, Li L, Tognin S, Wu Q, Pettersson-Yeo W, Lui S, Huang X, Marquand AF, Mechelli A. Using structural neuroanatomy to identify trauma survivors with and without post-traumatic stress disorder at the individual level. Psychol Med. 2014;44:195–203.

14. Zhang Q, Wu Q, Zhu H, He L, Huang H, Zhang J, Zhang W. Multimodal MRI-Based Classification of Trauma Survivors with and without Post-Traumatic Stress Disorder. Front Neurosci. 2016;10:292.

15. Yuan M, Qiu C, Meng Y, Ren Z, Yuan C, Li Y, Gao M, Lui S, Zhu H, Gong Q, Zhang W. Pre-treatment Resting-State Functional MR Imaging Predicts the Long-Term Clinical Outcome After Short-Term Paroxtine Treatment in Post-traumatic Stress Disorder. Front Psychiatry. 2018;9:532.

16. Etkin A, Maron-Katz A, Wu W, Fonzo GA, Huemer J, Vértes PE, Patenaude B, Richiardi J, Goodkind MS, Keller CJ, Ramos-Cejudo J, Zaiko YV, Peng KK, Shpigel E, Longwell P, Toll RT, Thompson A, Zack S, Gonzalez B, Edelstein R, Chen J, Akingbade I, Weiss E, Hart R, Mann S, Durkin K, Baete SH, Boada FE, Genfi A, Autea J, Newman J, Oathes DJ, Lindley SE, Abu-Amara D, Arnow BA, Crossley N, Hallmayer J, Fossati S, Rothbaum BO, Marmar CR, Bullmore ET, O’Hara R. Using fMRI connectivity to define a treatment-resistant form of post-traumatic stress disorder. 2019;11:eaal3236.

17. van Rooij SJ, Kennis M, Sjouwerman R, van den Heuvel MP, Kahn RS, Geuze E. Smaller hippocampal volume as a vulnerability factor for the persistence of post-traumatic stress disorder. Psychol Med. 2015;45:2737–2746.

18. Kennis M, van Rooij SJ, Tromp do PM, Fox AS, Rademaker AR, Kahn RS, Kalin NH, Geuze E. Treatment Outcome-Related White Matter Differences in Veterans with Posttraumatic Stress Disorder. Neuropsychopharmacology. 2015;40:2434–2442.

19. van Rooij SJ, Kennis M, Vink M, Geuze E. Predicting Treatment Outcome in PTSD: A Longitudinal Functional MRI Study on Trauma-Unrelated Emotional Processing. Neuropsychopharmacology. 2016;41:1156–1165.

20. Kennis M, Rademaker AR, van Rooij SJ, Kahn RS, Geuze E. Resting state functional connectivity of the anterior cingulate cortex in veterans with and without post-traumatic stress disorder. Hum Brain Mapp. 2015;36:99–109.

21. Kennis M, van Rooij SJ, van den Heuvel MP, Kahn RS, Geuze E. Functional network topology associated with posttraumatic stress disorder in veterans. Neuroimage Clin. 2016;10:302–309.

22. Blake DD, Weathers FW, Nagy LM, Kaloupek DG, Gusman FD, Charney DS, Keane TM. The development of a Clinician-Administered PTSD Scale. J Trauma Stress. 1995;8:75–90.

23. First MB, Spitzer RL, Gibbon M, Williams JB. Structured clinical interview for DSM-IV axis I disorders. New York: New York State Psychiatric Institute. 1995.

24. Brady K, Pearlstein T, Asnis GM, Baker D, Rothbaum B, Sikes CR, Farfel GM. Efficacy and safety of sertraline treatment of posttraumatic stress disorder: a randomized controlled trial. JAMA. 2000;283:1837–1844.

25. Davidson JR, Rothbaum BO, van der Kolk BA, Sikes CR, Farfel GM. Multicenter, double-blind comparison of sertraline and placebo in the treatment of posttraumatic stress disorder. Arch Gen Psychiatry. 2001;58:485–492.

26. World Medical A. World Medical Association Declaration of Helsinki: ethical principles for medical research involving human subjects. JAMA. 2013;310:2191–2194.

27. Pruim RHR, Mennes M, van Rooij D, Llera A, Buitelaar JK, Beckmann CF. ICA-AROMA: A robust ICA-based strategy for removing motion artifacts from fMRI data. Neuroimage. 2015;112:267–277.

28. Beckmann CF, Smith SM. Probabilistic independent component analysis for functional magnetic resonance imaging. IEEE Trans Med Imaging. 2004;23:137–152.

29. Biswal BB, Mennes M, Zuo XN, Gohel S, Kelly C, Smith SM, Beckmann CF, Adelstein JS, Buckner RL, Colcombe S, Dogonowski AM, Ernst M, Fair D, Hampson M, Hoptman MJ, Hyde JS, Kiviniemi VJ, Kotter R, Li SJ, Lin CP, Lowe MJ, Mackay C, Madden DJ, Madsen KH, Margulies DS, Mayberg HS, McMahon K, Monk CS, Mostofsky SH, Nagel BJ, Pekar JJ, Peltier SJ, Petersen SE, Riedl V, Rombouts SA, Rypma B, Schlaggar BL, Schmidt S, Seidler RD, Siegle GJ, Sorg C, Teng GJ, Veijola J, Villringer A, Walter M, Wang L, Weng XC, Whitfield-Gabrieli S, Williamson P, Windischberger C, Zang YF, Zhang HY, Castellanos FX, Milham MP. Toward discovery science of human brain function. Proc Natl Acad Sci U S A. 2010;107:4734–4739.

30. Smith SM, Nichols TE. Threshold-free cluster enhancement: addressing problems of smoothing, threshold dependence and localisation in cluster inference. Neuroimage. 2009;44:83–98.

31. Rasmussen CE, Williams CKI: Gaussian Processes for Machine Learning. Cambridge, Massachusetts, The MIT Press; 2006.

32. Ruan J, Bludau S, Palomero-Gallagher N, Caspers S, Mohlberg H, Eickhoff SB, Seitz RJ, Amunts K. Cytoarchitecture, probability maps, and functions of the human supplementary and pre-supplementary motor areas. Brain Struct Funct. 2018;223:4169–4186.

33. Eickhoff SB, Bzdok D, Laird AR, Roski C, Caspers S, Zilles K, Fox PT. Co-activation patterns distinguish cortical modules, their connectivity and functional differentiation. Neuroimage. 2011;57:938–949.

34. van Rooij SJH, Jovanovic T. Impaired inhibition as an intermediate phenotype for PTSD risk and treatment response. Prog Neuropsychopharmacol Biol Psychiatry. 2019;89:435–445.

35. van Waarde JA, Scholte HS, van Oudheusden LJ, Verwey B, Denys D, van Wingen GA. A functional MRI marker may predict the outcome of electroconvulsive therapy in severe and treatment-resistant depression. Mol Psychiatry. 2015;20:609–614.

36. Bludau S, Eickhoff SB, Mohlberg H, Caspers S, Laird AR, Fox PT, Schleicher A, Zilles K, Amunts K. Cytoarchitecture, probability maps and functions of the human frontal pole. Neuroimage. 2014;93 Pt 2:260–275.

37. Gilbert SJ, Spengler S, Simons JS, Steele JD, Lawrie SM, Frith CD, Burgess PW. Functional specialization within rostral prefrontal cortex (area 10): a meta-analysis. J Cogn Neurosci. 2006;18:932–948.

38. Burgess PW, Wu H-C: Rostral Prefrontal Cortex (Brodmann Area 10). in Principles of Frontal Lobe Function 2013. pp. 524–544.

39. Lane RD, Ryan L, Nadel L, Greenberg L. Memory reconsolidation, emotional arousal, and the process of change in psychotherapy: New insights from brain science. Behav Brain Sci. 2015;38:e1.

40. Colvonen PJ, Glassman LH, Crocker LD, Buttner MM, Orff H, Schiehser DM, Norman SB, Afari N. Pretreatment biomarkers predicting PTSD psychotherapy outcomes: A systematic review. Neurosci Biobehav Rev. 2017;75:140–156.

41. Ashburner J, Friston KJ. Unified segmentation. Neuroimage. 2005;26:839–851.

42. Ashburner J. A fast diffeomorphic image registration algorithm. Neuroimage. 2007;38:95–113.

43. Avants BB, Epstein CL, Grossman M, Gee JC. Symmetric diffeomorphic image registration with cross-correlation: evaluating automated labeling of elderly and neurodegenerative brain. Med Image Anal. 2008;12:26-

44. Jenkinson M, Beckmann CF, Behrens TE, Woolrich MW, Smith SM. Fsl. Neuroimage. 2012;62:782–790.

45. Tustison NJ, Avants BB, Cook PA, Zheng Y, Egan A, Yushkevich PA, Gee JC. N4ITK: improved N3 bias correction. IEEE Trans Med Imaging. 2010;29:1310–1320.

46. Zhang Y, Brady M, Smith S. Segmentation of brain MR images through a hidden Markov random field model and the expectation-maximization algorithm. IEEE Trans Med Imaging. 2001;20:45–57.

47. Greve DN, Fischl B. Accurate and robust brain image alignment using boundary-based registration. Neuroimage. 2009;48:63–72.

48. Ciric R, Wolf DH, Power JD, Roalf DR, Baum GL, Ruparel K, Shinohara RT, Elliott MA, Eickhoff SB, Davatzikos C, Gur RC, Gur RE, Bassett DS, Satterthwaite TD. Benchmarking of participant-level confound regression strategies for the control of motion artifact in studies of functional connectivity. Neuroimage. 2017;154:174–187.

49. Jenkinson M, Bannister P, Brady M, Smith S. Improved optimization for the robust and accurate linear registration and motion correction of brain images. Neuroimage. 2002;17:825–841.

50. Van Dijk KR, Hedden T, Venkataraman A, Evans KC, Lazar SW, Buckner RL. Intrinsic functional connectivity as a tool for human connectomics: theory, properties, and optimization. J Neurophysiol. 2010;103:297–321.

51. Abou Elseoud A, Littow H, Remes J, Starck T, Nikkinen J, Nissila J, Timonen M, Tervonen O, Kiviniemi V. Group-ICA Model Order Highlights Patterns of Functional Brain Connectivity. Front Syst Neurosci. 2011;5:37.

52. Cerliani L, Mennes M, Thomas RM, Di Martino A, Thioux M, Keysers C. Increased Functional Connectivity Between Subcortical and Cortical Resting-State Networks in Autism Spectrum Disorder. JAMA Psychiatry. 2015;72:767–777.

53. Beckmann CF, Mackay CE, Filippini N, Smith SM. Group comparison of resting-state FMRI data using multi-subject ICA and dual regression. Neuroimage. 2009;47:S148.

54. Marquand A, Howard M, Brammer M, Chu C, Coen S, Mourao-Miranda J. Quantitative prediction of subjective pain intensity from whole-brain fMRI data using Gaussian processes. Neuroimage. 2010;49:2178–2189.

55. Minka TP: A family of algorithms for approximate Bayesian inference. Massachusetts Institute of Technology; 2001.

